# Divide-and-conquer quantum algorithm for hybrid *de novo* genome assembly of short and long reads

**DOI:** 10.1101/2023.09.19.558544

**Authors:** Jing-Kai Fang, Yue-Feng Lin, Jun-Han Huang, Yibo Chen, Gao-Ming Fan, Yuhui Sun, Guanru Feng, Cong Guo, Tiejun Meng, Yong Zhang, Xun Xu, Jingen Xiang, Yuxiang Li

**Author notes:** These authors contributed equally to this work (Jing-Kai Fang, Yue-Feng Lin, Jun-Han Huang).

## Abstract

Computational biology holds immense promise as a domain that can leverage quantum advantages due to its involvement in a wide range of challenging computational tasks. Researchers have recently explored the applications of quantum computing in genome assembly implementation. However, the issue of repetitive sequences remains unresolved. In this paper, we propose a hybrid assembly quantum algorithm using high-accuracy short reads and error-prone long reads to deal with sequencing errors and repetitive sequences. The proposed algorithm builds upon the variational quantum eigensolver and utilizes divide-and-conquer strategies to approximate the ground state of larger Hamiltonian while conserving quantum resources. Using simulations of 10-qubit quantum computers, we address problems as large as 140 qubits, yielding optimal assembly results. The convergence speed is significantly improved via the problem-inspired ansatz based on the known information about the assembly problem. Besides, entanglement within quantum circuits is qualitatively verified to notably accelerate the assembly path optimization.

## 1 Introduction

DNA sequencing technology has dramatically transformed the fields of biology and medicine in the past few decades. This revolutionary tool allows researchers to decode the genetic blueprints of living organisms, leading to breakthroughs such as early cancer diagnosis and detection of inherited diseases. The throughput and speed of DNA sequencing have increased exponentially over the years, surpassing even the Moore’s law that predicts the growth of computational power [1]. The Sanger sequencing platform, back in 1987, was only capable of sequencing approximately 1000 nucleotides per day [1–3]. This limitation rendered the sequencing and assembly of the complete human genome by the Human Genome Project a 13-year endeavor. In contrast, present technologies allow for the sequencing of an entire human genome within mere hours. However, sequencing is just one facet of the challenge. Genome reconstruction is an indispensable subsequent step to pave the way for comprehensive studies. With the rapid developing of sequencing technology, there has been a corresponding surge in data volume that amplifies computational demands, prompting the evolution of diverse assembly algorithms.

One of the early assembly algorithms is the overlaplayout-consensus (OLC) algorithm [4–8]. The OLC algorithm transforms the genome assembly problem into a graph problem where each vertex represents a read and edges represent overlaps among all reads, aiming to find a Hamiltonian path in the graph, which is equivalent to the traveling salesman problem, a famous NP-hard combinatorial optimization problem [9]. The OLC approach has been widely used in various genome assemblers for long DNA fragments. Another different algorithm, based on de Bruijn graphs, has been applied in a series of assemblers, called Eulerian assemblers [10], such as SOAPdenovo [11], Spades [12] and ABySS [13]. In this algorithm, vertices in the graph no longer represent reads, but rather k-mers generated from the reads, and the assembly process can be reformulated as an Eulerian path problem. Unlike the intractable complexity of OLC, the de Bruijn graph assemblers are suitable for the assembly based on next-generation sequencing data that produces a vast number of high-accuracy short reads.

The next-generation sequencing has been extensively used due to its accuracy and low cost. Nevertheless, the limited length of short reads makes it difficult to complete the assembly of highly repetitive complex regions in genomes. The advent of the third-generation sequencing platform such as Oxford Nanopore Technology (ONT) [14] and Pacific Biosciences (PacBio) [15–17] has enabled the generation of long reads with a length of more than 10kbp, which can span over multiple genomic repeats and improve the contiguity of assembly. However, these long reads also have a high error rate and are not cost-friendly, limiting their applicability for large-scale genome projects.

To overcome these challenges, hybrid assembly approaches combining short and long reads have been developed. These methods have been shown to bring unique benefits by complementing the strengths and weaknesses of each read type. For example, Koren *et al*. [18] and Au *et al*. [19] applied short reads to correct long reads and then assembled the corrected long reads. These approaches require high coverage and usually huge computing resources. Deshpande *et al*. [20], Antipov *et al*. [21] and Wick *et al*. [22] used another approach that assembled the short reads first to generate precise contigs and employed long reads for scaffolding. The hybrid assembly provides an alternative cost-effective way since it requires fewer long reads than long-read-only methods. Some recent studies [23, 24] have shown that hybrid assembly is superior to longread-only methods in terms of correctness, contiguity and completeness. Nevertheless, hybrid assembly is still computationally demanding, requiring powerful computational resources and storage capacity, especially for large and complex genomes.

Whereas, concerns have been raised regarding the ability of supercomputers to handle the explosive growth of sequencing data, particularly as classical computers are already approaching their physical limits. Given these limitations of classical computing, it is crucial to explore alternative computational paradigms. Quantum computers promise a new solution to this challenge. Especially in terms of enhanced computational efficiency and capability for handling complex optimization problems [25, 26]. Numerous quantum algorithms have been proposed and demonstrated promising quantum advantages. Some of them have been applied to various bioinformatics problems, such as *de novo* assembly [27– 29], sequence alignment [30, 31], protein folding [32–34], and phylogenetic tree inference [35], which inspired us to achieve hybrid assembly using quantum computing.

The previous studies [27–29] proposed several OLC-based assembly quantum algorithms, including quantum annealing and variational quantum algorithms, and provided a proof of the application of quantum computing to the assembly problem, where sequencing error was also taken into account in Ref. [29]. However, there are inevitable repeats in DNA sequences, and they cannot be solved before assembly. The classical assemblers solve the problem by indexing reads near the repeats and seeking out the reads that contain the entire repetitive region. But it is worth mentioning that none of quantum algorithms have addressed the issue of repetitive sequences.

In this paper, we propose a quantum algorithm for hybrid *de novo* genome assembly using short and long reads. In our algorithm, short reads and long reads are utilized to deal with the assembly path conflicts caused by sequencing errors or repetitive sequences. Concretely, we first map the hybrid *de novo* assembly problem with repeats and sequencing errors onto a combinatorial optimization problem. Then we utilize the variational quantum eigensolver (VQE) framework and introduce the divide-and-conquer strategy to obtain optimal assembly sequences with fewer quantum resources in the noisy intermediate-scale quantum (NISQ) era, characterized by quantum devices that are susceptible to errors but can still provide computational advantages [36]. Furthermore, a problem-inspired ansatz is proposed to significantly improve the hybrid assembly quantum algorithm.

## 2 Results

### 2.1 Hybrid assembly quantum algorithm

The workflow of the hybrid assembly quantum algorithm is presented in Figure 1. The sequencing data of short reads and long reads are processed on quantum computers or quantum simulators after a encoding step. This generates a state that can be decoded to obtain the assembly sequences. The details are shown below.

**Fig. 1.**
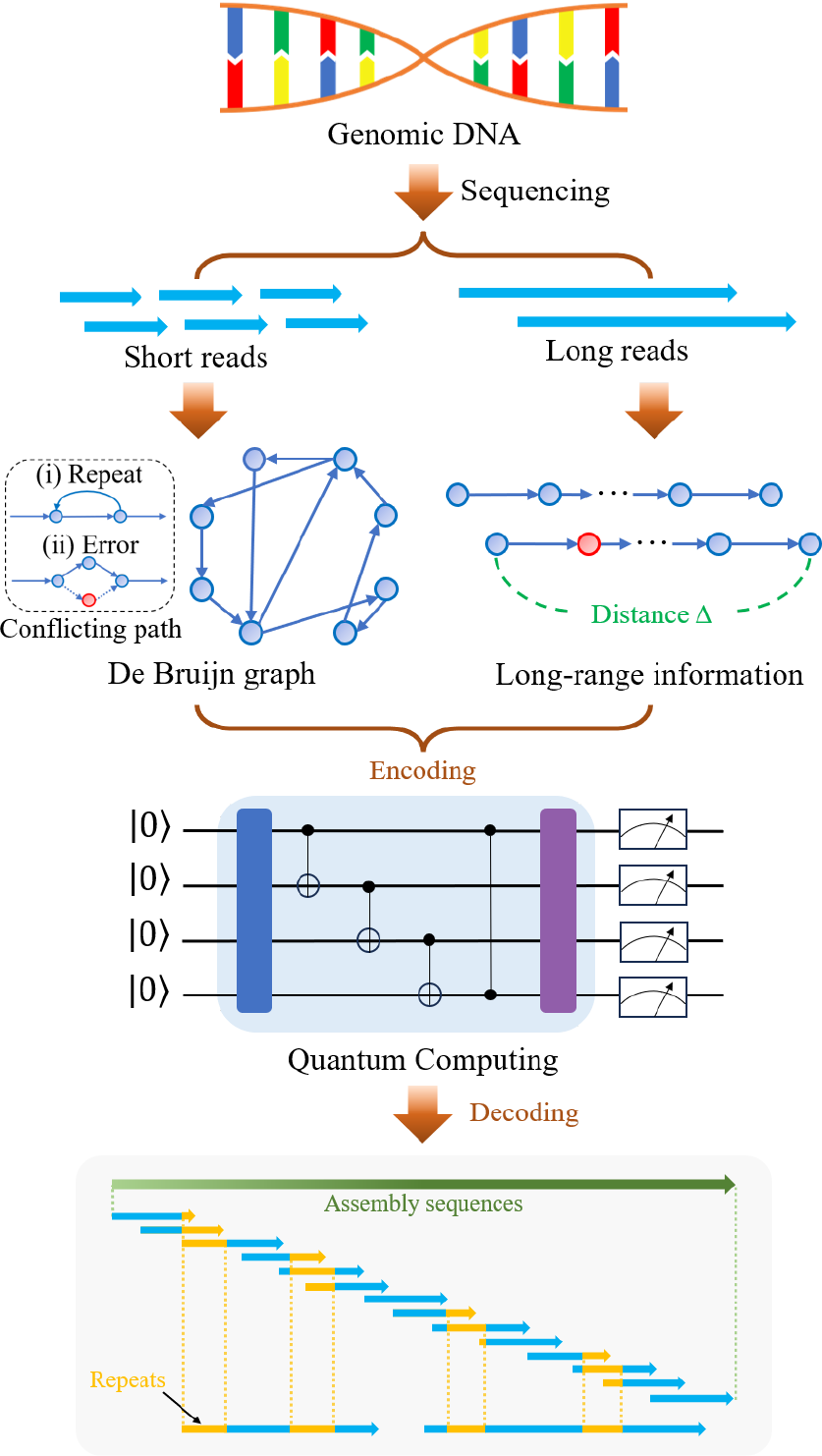
Schematic overview of the quantum algorithm for hybrid *de novo* genome assembly. The short reads are used to construct the de Bruijn graphs, while the long reads are used to provide longrange linking between nodes. Here, the nodes represent k-mers and the red ones mean low-frequency k-mers from sequencing errors. Connections between nodes are based on the common k−1 suffix and prefix. Inevitable path conflicts are caused by sequencing errors or repetitive sequences. The short-read and long-read data are encoded together into a optimization model solvable on a quantum device. The results are then decoded to obtain the assembly sequences.

We first propose a mapping model of the hybrid *de novo* assembly problem involving repetitive sequences and sequencing errors. Due to the path conflicts brought by repetitive sequences and sequencing errors, it is not feasible to directly determine an Eulerian path within the de Bruijn graph [10] that faithfully reconstructs the original sequences. To overcome this obstacle, we introduce long-reads data of third-generation sequencing and encode the information into the weight of the cost function. Then we can construct the cost function of hybrid *de novo* assembly problem.

First, we need to encode the de Bruijn graph. Let binary variables 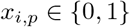 be the occupation number of every node at each position, where *i* ∈ {1, 2, …,n} represents i-th node and *p* ∈ {1, 2, …, p_max_} represents the position in the assembly path. 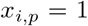 means that i-th node is chosen at position p in the path. The possible assembly path can be expressed as a quantum state 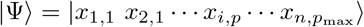. Note that for a graph with n nodes, the maximum of p must be larger than n if there are repeats in DNA sequences. Conservatively, we can take p_max_ = 2n. Then the cost function of the hybrid *de novo* assembly problem is formulated as

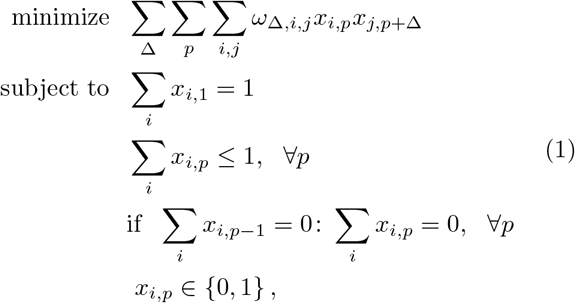

where the ω_Δ,*i,j*_ is the weight of *k*-mer *i* and *j* separated by an inter-nucleotide distance Δ. For the weight of two adjacent nodes, we set ω_1,*i,j*_ = −α to indicate that two k-mers are correctly sequenced while setting ω_1,*i,j*_ =− β if the frequency of k-mers is less than 10% (consider as sequencing errors), where 0 < β ≪ α. For nodes that are not connected, a large positive weight γ is given as a penalty. Additionally, by introducing the long-read data from third-generation sequencing as shown in Figure 1, we can determine whether any two nodes are separated by a distance Δ. The long-range links between the nodes depend on the length of long-read sequences that generally exceed 10 kbp. If two nodes are identified with a distance of Δ in long read, we assign the corresponding term with a negative weight ω_Δ,*i,j*_ = −η to contribute to the cost function. Otherwise we set it to be zero. In order to solve the problem of repetitive sequences, the distance Δ must be greater than the sum of the lengths of the repeats and the absolute value of ω_Δ,*i,j*_ should be as large as possible to against the total weight of the cyclic paths caused by repetitive sequences.

Next, we construct the Hamiltonian of the hybrid *de novo* assembly problem according to the cost function in Equation (1). The objective term can be directly written as

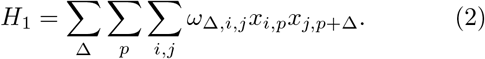

Then we consider the constraints of the cost function. The first constraint term of Equation (1) ensures that the first position (*p* = 1) of the assembly path must be one and only one node. Thus, the Hamiltonian of this constraint term can be formulated as

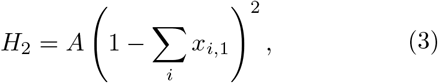

where coefficient *A* is the penalty parameter. The second and third constraint terms ensure the contiguity of the assembly sequence, that is, the assembly path is unique and continuous. Mathematically, for all *p*, the term 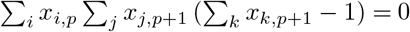 ensures an unforked path, and the term 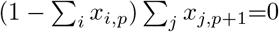 prevents the path from being discontinuous. Thus, the Hamiltonian of the second and third constraint terms can be combined as

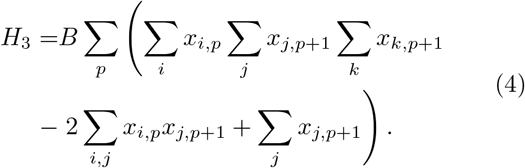

Here, the parameters must satisfy 0 < B < A to ensure that the contribution of all penalty terms is not less than zero. Finally, the whole Hamiltonian of the hybrid *de novo* assembly problem is expressed as

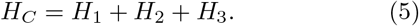

Thus, the optimal assembly path with cycle caused by repeats in the de Bruijn graph can be effectively represented as a continuous and unforked path. According to this Hamiltonian, parameterized quantum circuits are constructed and applied in the VQE approach to calculate the minimum eigenvalue and the corresponding eigenvector of H_*C*_ that exactly matches the optimal assembly path (Methods). Eventually, a decoding process is added to yield the desired assembly sequence as shown in Figure 1.

### 2.2 Numerical simulations

The proposed hybrid assembly quantum algorithm is implemented in SpinQit with Torch simulator. We analyze the convergence properties of various quantum circuits, including two distinct types of hardware-efficient ansatzes (HEA), as well as a problem-inspired ansatz based on the known information about the assembly problem. The experiments are conducted on Dell equipped with 2.90 GHz Intel Core i7-10700 and 16.0 GB memory, and operated under the Windows 11 operating system.

#### Toy model

We illustrate hybrid assembly quantum algorithm for the toy model as shown in Figure 2. The nodes in the toy model represent k-mers or a group of k-mers generated by splitting short reads. Note that we treat three consecutive k-mers as a single node in the de Bruijn graph to further simplify the model. And long-read data is introduced as weights ω_Δ,*i,j*_ to aid assembly.

**Fig. 2.**
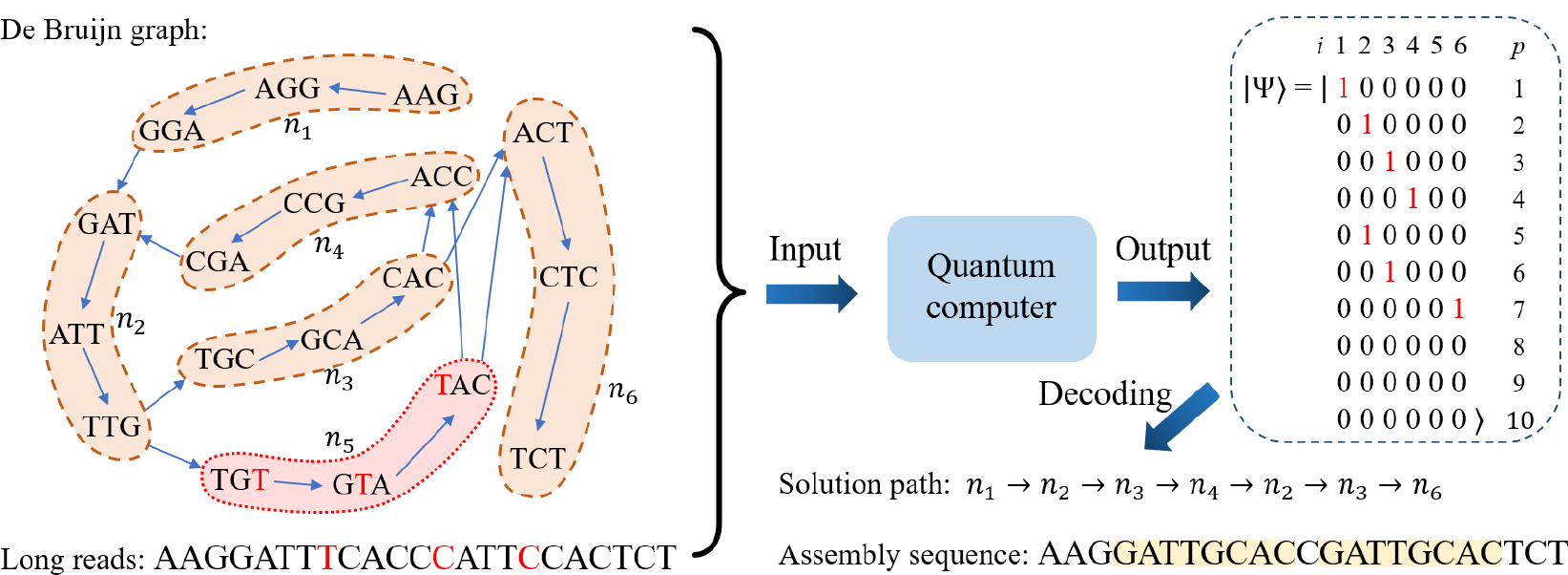
Schematic diagram of the toy model. Both short-read and long-read sequencing data are used, where the red characters represent sequencing errors. To optimize qubit utilization and streamline the model, a strategy is implemented where three consecutive k-mers are consolidated into a single node. And the nodes in the red dotted circle are low-frequency k-mers from sequencing errors. The connection of the nodes is established based on the shared k− 1 suffix and prefix. The nodes n_2_ and n_3_ represent repeat regions, and the node n_5_ represents sequencing errors. The toy model is input into a quantum computer for simulation, resulting in the generation of the final quantum state of the binary string |Ψ⟩. The optimal assembly sequence is then obtained through the process of decoding the binary string.

In this toy model, we assume that the correct assembly path is n_1_ → n_2_ → n_3_ → n_4_ → n_2_ → n_3_ → n_6_. Here, we set p_max_ = 10 to allow more wrong assembly solution to verify the reliability of our approach. Moreover, we utilize the long-range linking information to ensure that the loop n_2_→ n_3_→ n_4_→ n_2_ does not occur twice by setting α = 2, β = 1, γ = 20, η = 10, respectively. Considering the quantum resources required is quadratic to the problem size, a divide-and-conquer strategy is designed in the VQE framework (Methods). Thus, the VQE algorithm is used to obtain optimal assembly result by simulations of 6-qubit quantum computers. In this case, the exact loss value is − 27 and it is verified that the corresponding assembly sequence AAGGATTGCACCGATTGCACTCT is correctly extracted.

#### Entangling capability

In practice, the final results of the hybrid assembly are significantly rely on the expressibility and entangling capability of ansatzes. Hardware-efficient ansatzes [37] are one of the widely used parameterized quantum circuits in the VQE algorithm, as they are designed based on several factors of the near-term quantum devices, such as qubit connectivity and restricted gate sets. Normally, it is constructed by the single-qubit gates with parameters that can be optimized, including *R*_*x*_, *R*_*y*_ and *R*_*z*_ gates, as well as two-qubit entangling gates, such as CNOT, CZ and iSwap gates.

In this work, we simulate the Hamiltonian of the toy model by the VQE approach with two different hardware-efficient ansatzes. Our goal is to examine the performance of optimization convergence with and without quantum entanglement. In Figures 3(a) and 3(b), the former called as HEA1 consists of *R*_*y*_ gate and *R*_*z*_ gate and CZ gate. The latter called as HEA2 is constructed with single-qubit rotations *R*_*y*_ and *R*_*z*_ only. In the experiment, we set the depth of quantum circuit *D*_*e*_ to be 2, and randomly generate 100 sets of initial parameters for HEA1 and HEA2. As illustrated in Figures 4(a) and 4(b), our findings indicate a significant correlation between the initial parameters and the convergence speed of various ansatzes. More importantly, we show that the average convergence speed of HEA1 is faster than that of HEA2. In Figure 4(c), the average iteration number for the convergence of HEA1 is 348, while it is 393 for HEA2. This means that the entangling capability of ansatzes can accelerate the convergence of the hybrid *de novo* assembly quantum algorithm using short and long reads.

**Fig. 3.**
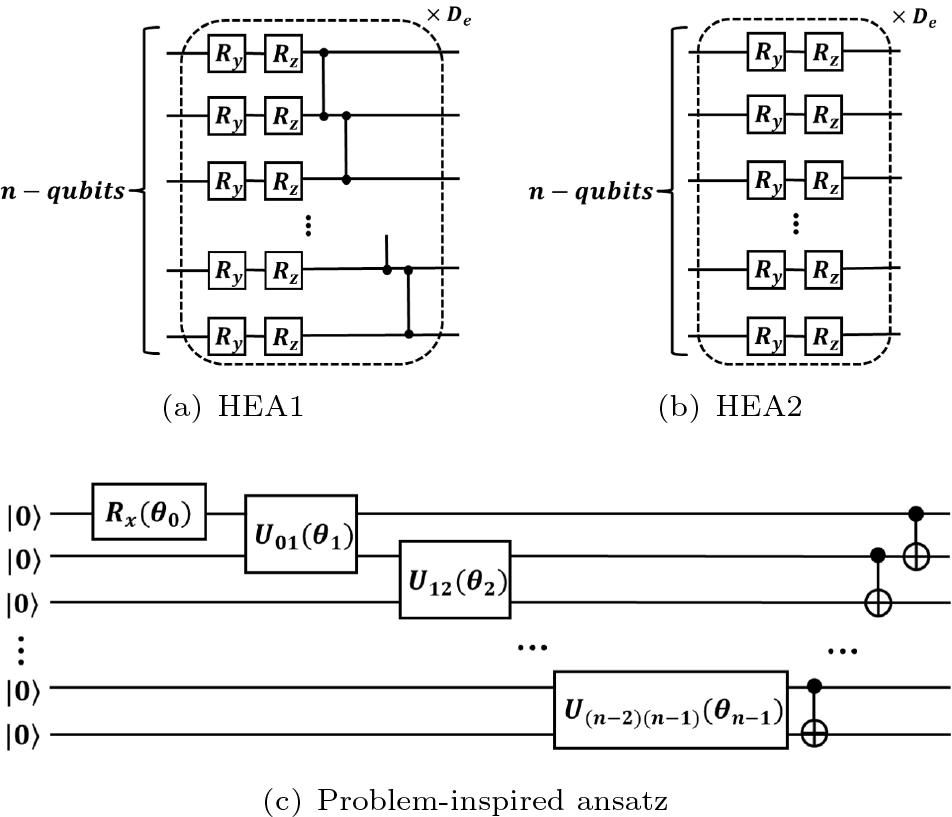
Variational ansatzes for the VQE algorithm. (a) HEA1 implemented by *Ry, Rz* and CZ gates. (b) HEA2 implemented by *Ry* and *Rz* gates. The quantum circuits in the dashed boxes repeat *De* times. (c) The problem-inspired ansatz based on the assembly problem (Methods).

**Fig. 4.**
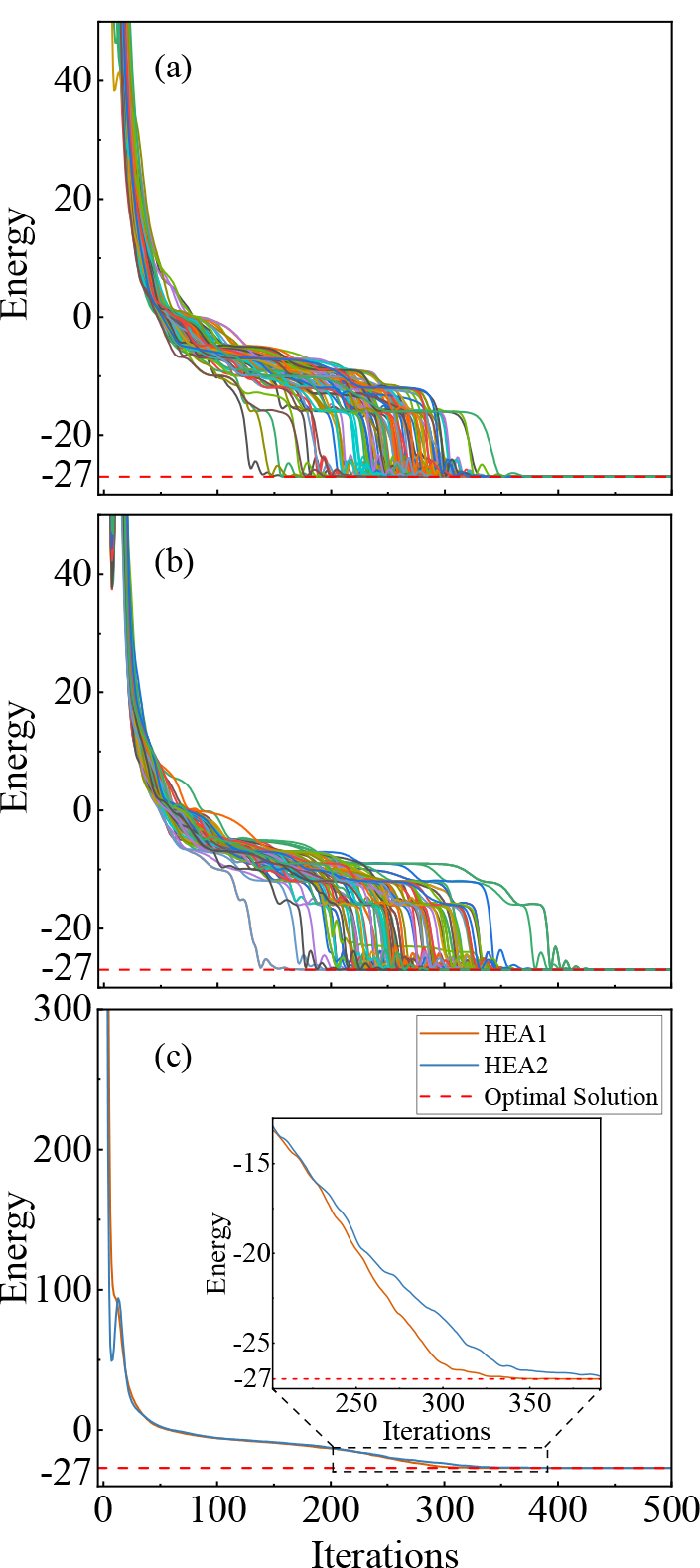
Simulation results of HEA. (a) and (b) show the convergence of HEA1 and HEA2 with 100 sets of random initial parameters, respectively. (c) The comparison of the average convergence between HEA1 and HEA2.

#### Problem-inspired ansatz

Unlike HEA, the probleminspired ansatz is designed with unique attributes of assembly problems. In the divide-and-conquer quantum algorithm, because there is at most one variable of 1 in the binary string 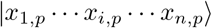, the t rial wavefunction of n qubits can be restricted to|ψ⟩ = Σ_*k*_ λ_*k*_ |k⟩, *k* ∈ {0, 2^0^, 2^1^,, 2^*n−*1^ }, where n is the number of nodes in de Bruijn graphs. The problem-inspired ansatz is constructed to prepare the candidate n-qubit trial wavefunction |ψ⟩, and more details can be found in Figure 3(c) and Methods.

Same as in the HEA case, we also randomly generate 100 sets of initial parameters for proposed problem-inspired ansatz. As shown in Figure 5(a), we show that the convergence speed of the problem-inspired ansatz is significantly faster than that of HEA. Furthermore, the average iteration number required for the convergence of the problem-inspired ansatz is 168 in Figure 5(b). The results verify that the problem-inspired ansatz can accelerate the convergence speed of the hybrid assembly quantum algorithm due to a reduced search space and fewer optimization parameters. Table 1 presents the quantum resources required for the three quantum circuits. In contrast to HEA, the problem-inspired ansatz employs a fixed circuit depth, yielding a streamlined computational process and fewer quantum gate requirements.

**Table 1.**
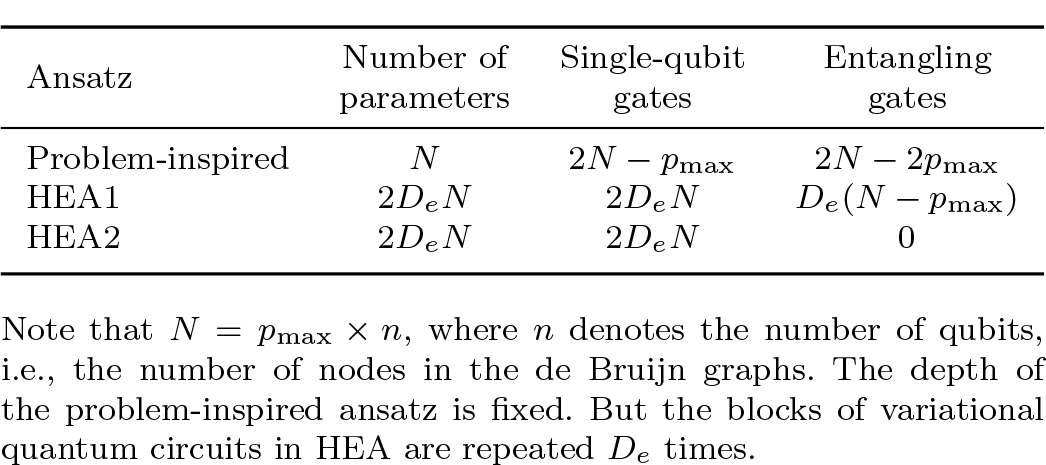
Cost estimates for the problem-inspired ansatz and HEA.

**Fig. 5.**
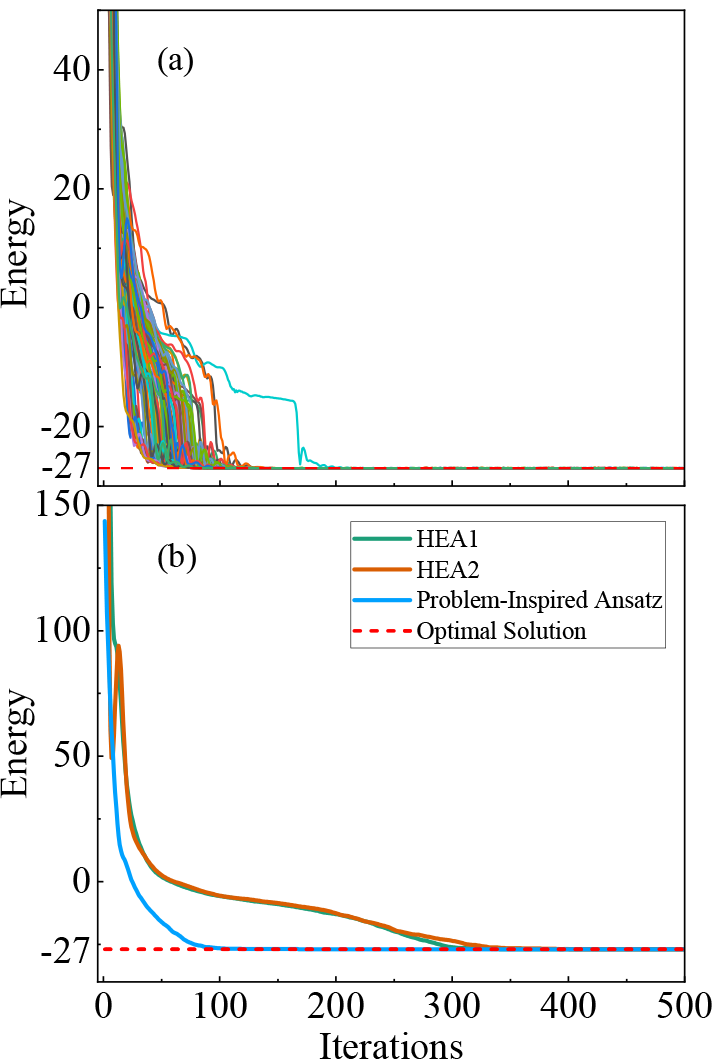
Simulation results of the problem-inspired ansatz. (a) The evaluation value for 100 groups of random initial parameters with the problem-inspired ansatz. (b) The comparison of the average convergence of HEA1, HEA2 and the problem-inspired ansatz.

#### Problem sizes of hundreds of qubits

Further, we evaluate the performance of problem-inspired ansatz on larger problems. Five different sizes of assembly problems, ranging from 6 nodes to 10 nodes, are illustrated in Figure 6 with number of qubits required for the corresponding Hamiltonian are 60, 77, 96, 117 and 140, respectively. Note that a divide-and-conquer strategy is employed in the VQE framework in order to calculate on quantum simulators. This approach enables more efficient calculations on quantum simulators by breaking down larger problems into more manageable sub-problems. Besides, we restrict the initial parameters of the problem-inspired ansatz to the range [0.3π, 0.5π], as opposed to the more common range of [0, π], This restriction led to a significant improvement in both the convergence speed and success rate of the optimization process. As evidenced by Figure 7(a), the ground-state energy of each Hamiltonian and the assembly path for each problem size are correctly optimized, serving as a validation of the problem-inspired ansatz on larger problems.

**Fig. 6.**
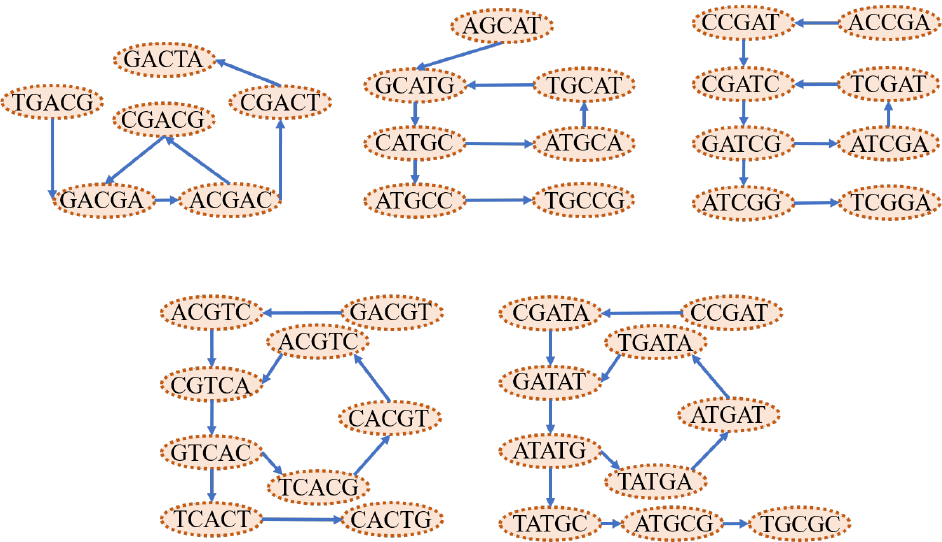
De Bruijn graphs for five different sizes of assembly problems, from 6 nodes to 10 nodes, are calculated by the probleminspired ansatz.

**Fig. 7.**
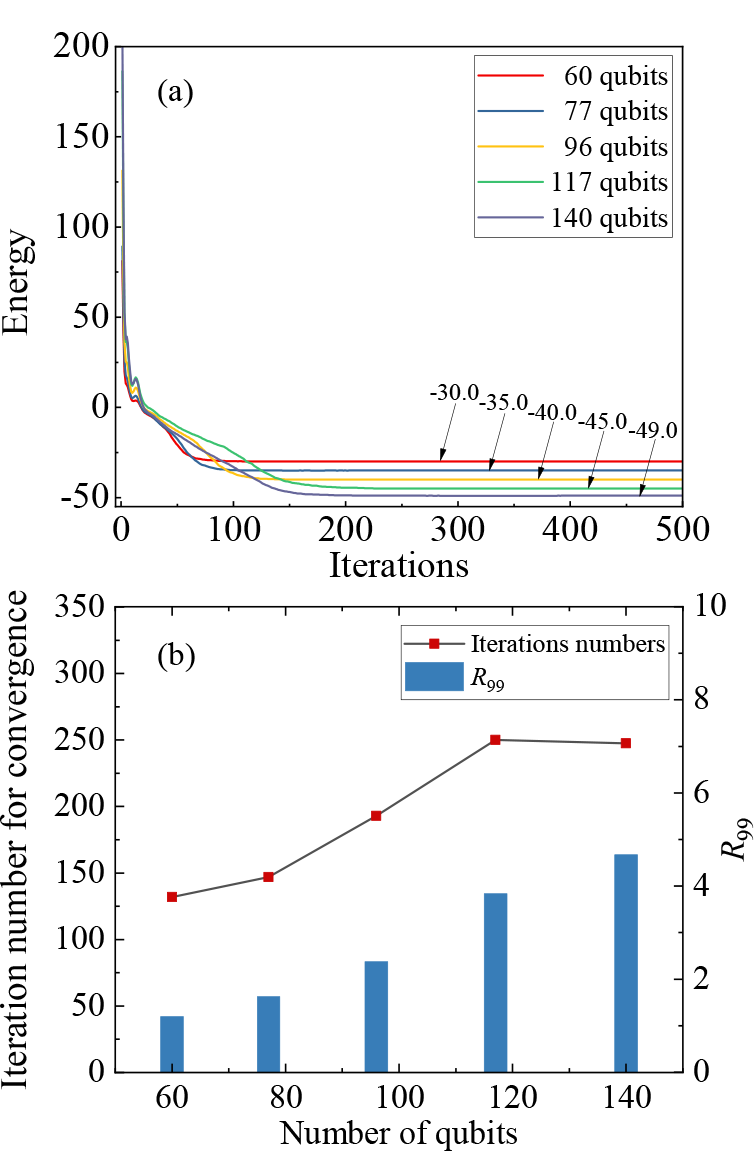
Simulation results of the problem-inspired ansatz for five different problem sizes. (a) The average convergence of different problem sizes. (b) The iteration number for convergence and the number of experiments required to find the optimal assembly results with a probability of 99%, called *R*_99_.

Finally, we measure the impact of problem sizes on computational cost by evaluating the average iteration number for convergence and *R*_99_ [27] of the five different sizes. Here, the parameter *R*_99_ represents the number of experiments required to find the optimal assembly sequence (ground state) with a probability of 99%. The algebraic expression of *R*_99_ is defined as

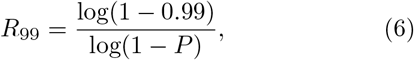

where *P* is the estimated success probability of each simulation. As shown in Figure 7(b), both the average iteration number for convergence and *R*_99_ increase linearly with the size of the problem. In addition, the computational cost is directly proportional to the number of qubits involved in the hybrid *de novo* assembly problem. This suggests that the proposed ansatz scales reasonably well for larger problem sizes.

## 3 Discussion

In this work, we propose a divide-and-conquer quantum algorithm for hybrid *de novo* genome assembly of short and long reads. We demonstrate that the optimized assembly sequences can be obtained by decoding quantum computing results for the ground states based on the VQE framework. Furthermore, we adopt divide-and-conquer strategies to extract the optimal assembly results with problem sizes up to hundreds of qubits, by employing simulations on few-qubit quantum computers.

The hybrid assembly quantum algorithm offers several advantages. Firstly, it integrates short reads from next-generation sequencing technology and long reads from third-generation sequencing technology to address assembly path conflicts. This hybrid sequencing strategy can leverage the complementary strengths of both sequencing platforms to improve the correctness and completeness of genome assembly and annotation. Secondly, it has the potential to significantly reduce the computational expense of simulating intricate quantum systems, which could enable practical applications of quantum computing in life sciences during the NISQ era. The *de novo* genome assembly problem is solved by mapping the DNA sequences into graph models. However, encoding a binary variable with multidimensional information requires a number of qubits that scales as O(n^2^), where n denotes the number of nodes in de Bruijn graphs. This presents a major challenge in optimizing assembly sequences when using the original VQE without any preprocessing, especially on NISQ devices. Consequently, in light of quantum resources, the proposed algorithm utilizes a divide-and-conquer strategy to alleviate the requirements of logical qubits, that is, to scale as O(n). In addition, our results show that quantum entanglement can be leveraged to expedite the optimization of genome assembly problem. Furthermore, our proposed problem-inspired ansatz significantly accelerates the optimization of assembly paths in the hybrid assembly quantum algorithm.

However, some issues may emerge in practical applications of the hybrid assembly quantum algorithm and deserve further investigation. Above all, a considerable number of qubits are required in practical assembly tasks even with divide-and-conquer strategies, which poses a huge challenge for near-term quantum devices. As the problem size increases, the parameter optimization of variational quantum algorithms becomes unpredictable and intractable. Fortunately, we find that narrowing the range of initial parameters can significantly improve the parameter optimization and the probability of obtaining the optimal assembly results. This also provides insights into solving other combinatorial optimization problems that are prone to local minima or barren plateaus as the number of qubits increases. In the near future, improving variational quantum algorithms and exploring quantum advantages will be primary goals in the field of quantum computing applications, especially in bioinformatics.

## 4 Methods

### 4.1 Variational quantum eigensolver

VQE is a hybrid quantum-classical algorithm for solving approximate minimum eigenvalues and eigenvectors of Hamiltonian. It relies on the Rayleigh-Ritz variational principle as follows:

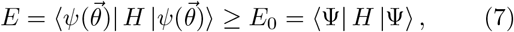

where 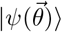 is the trial wavefunction prepared to extract the optimal assembly result by applying a parameterized quantum circuit to the initial state |ψ_0_ ⟩ . E is the expectation value of the Hamiltonian with 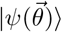. E_0_ and |Ψ ⟩ are the exact minimum eigenvalue and the corresponding eigenvector of the Hamiltonian H, respectively.

As shown in Figure 8, the procedure of VQE is implemented with two devices, quantum and classical computers. Typically, quantum computers are used to implement parameterized quantum circuits constructed by a set of quantum gates with adjustable parameters in order to prepare the trial wavefunction and measure the corresponding expectation value of the Hamiltonian. Meanwhile, the classical computer optimizes the parameters of quantum gates and transmits them to the quantum computer to initiate the next iteration. This iterative process continues until a predefined convergence criteria is satisfied. Explicitly, we introduce the key components of the VQE algorithm below.

**Fig. 8.**
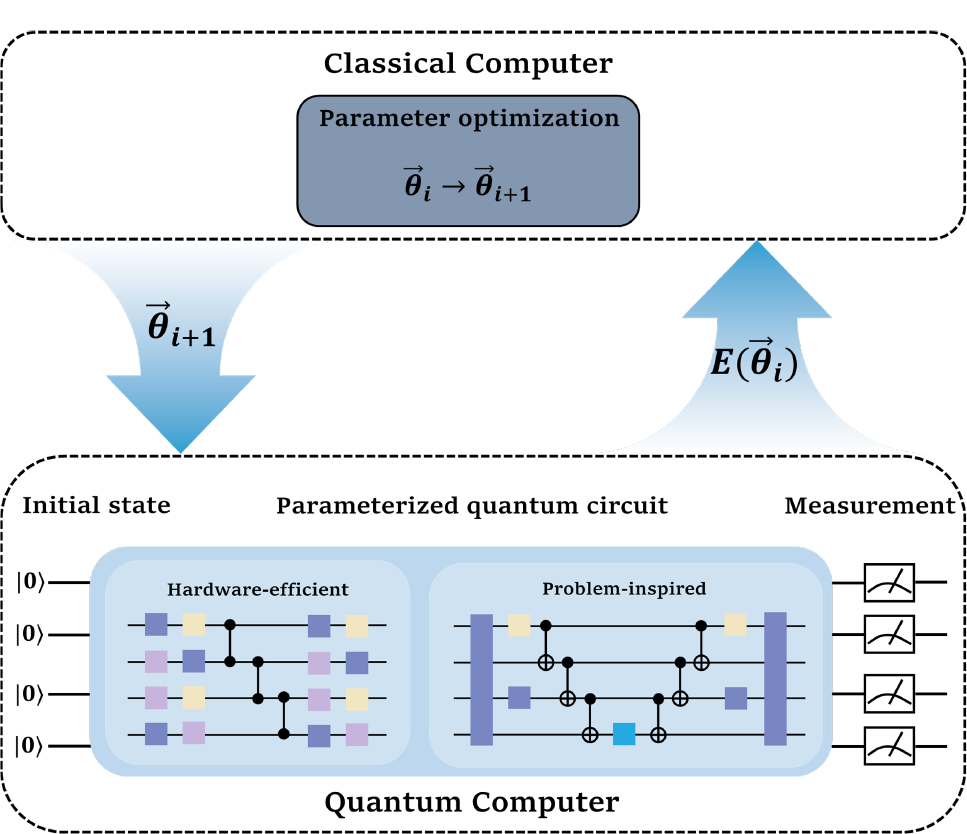
Scheme of the VQE method. VQE is a hybrid quantum-classical algorithm, where quantum processing units are used to prepare the trial wavefunction by parameterized quantum circuits and measure the corresponding expectation value of the Hamiltonian, and classical processing units are used to update the parameters.

#### Hamiltonian construction

For VQE, the first step is to construct the Hamiltonian for describing the various attributes of the systems, which are in general chemical molecules [38], combinatorial optimization problems [39], many-body physics [40] and so on. Since the quantum computers only measure observables that can be expressed as linear combinations of Pauli operators { I, X, Y, Z }, the Hamiltonian needs to be transformed to the corresponding terms.

#### Parameterized quantum circuits

The parameterized quantum circuit, also known as ansatz, is designed to prepare the trial wavefunction 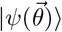 . Undeniably, since the VQE was first proposed by Peruzzo *et al*. in 2014 [41], the research on constructing a good ansatz for specific problems has attracted considerable interest. Now, various forms of parameterized quantum circuits have been proposed, such as hardware-efficient ansatzes [37], problem-inspired ansatzes [38, 39, 42–47] and variable-structure ansatzes [48, 49]. Generally speaking, a good ansatz includes the following two attributes: (i) the sufficient expressibility ensures that the trial wavefunction contains the ground state wavefunction, and (ii) the parameter search space is reduced as much as possible to speed up the convergence of the VQE algorithm. In our work, we design a problem-inspired ansatz to reduce the search space and boost the converging process for hybrid *de novo* genome assembly.

#### Parameter optimization

After transforming the Hamiltonian and preparing the trial wavefunction, the expectation value of the Hamiltonian can be measured by quantum computer. The measurement results are subsequently transmitted to the classical computer to update the parameters in the quantum circuits for the next iteration. Parameter optimization is necessary for any variational algorithms to obtain an approximate solution. The parameters of quantum circuits will be updated iteratively with some efficient optimization strategies, such as gradient descent methods [50, 51], as well as gradient-free methods [52, 53], until convergence.

### 4.2 Implementation

As mentioned above, the value of binary variable x_*i,p*_ indicates whether or not to choose the node *i* at the position p in the assembly path. And the possible assembly path can be expressed as a quantum state 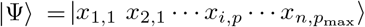 .

#### Qubit mapping

To construct the qubit Hamiltonian of the hybrid *de novo* assembly problem from Equation (5), the common approach is to map the binary variable *x*_*i,p*_ to Pauli *Z* and identity matrix *I* as follow:

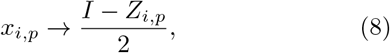

where *Z*_*i,p*_ is a Pauli *Z* acting on the [p(n− 1) + i]-th qubit. Correspondingly, Equation (5) can be rewritten as

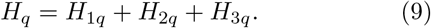

It is apparent that the dimension of the Hamiltonian system represented by H_*q*_ is 2^*N*^, where N = np_max_ denotes the required number of qubits.

In the NISQ era, it is challenging to employ the original VQE algorithm to simulate and estimate the approximate minimum eigenvalue of the Hamiltonian *H*_*q*_ without any preprocessing. Consequently, in light of the original VQE algorithm, we propose a divideand-conquer strategy to alleviate the requirements of logical qubits. To be specific, we leverage Hamiltonian decomposition and state decomposition to estimate the approximate ground-state energy of a *N*-qubit Hamiltonian on p_max_ quantum processing units with n qubits.

#### Hamiltonian decomposition

Once qubit mapping is performed, the Hamiltonian *H*_*q*_ can be expressed in the following form:

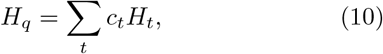

where *H*_*t*_ denotes a component of the Hamiltonian *H*_*q*_. *H*_*t*_ can be constructed using tensor products of the Pauli *Z* operator and the identity operator I. Its dimension is 2^*N*^ . We further decompose *H*_*t*_ into p_max_ Hamiltonians denoted as 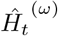,

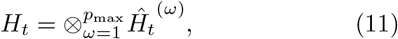

where the dimension of each Hamiltonian 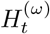 is 2^*n*^ Therefore, the Hamiltonian *H*_*q*_ can be rewritten as

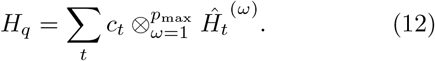

#### State decomposition

The quantum state of assembly path can be expressed using the tensor product notation,

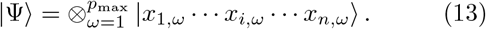

In VQE, each quantum state |x_1,*ω*_ …x_*i,ω*_ …x_*n,ω*_⟩ can be obtained from a trial wavefunction |*ψ*^*(ω*)^(*θ*)⟩

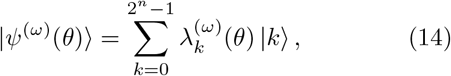

where |k⟩ are the n-qubit computational bases of |ψ^(*ω*)^(θ)⟩ and the parameters 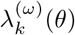 satisfy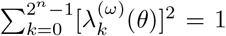 In general, the optimal assembly path corresponds to a non-superposed quantum state.

Therefore, the trial wavefunction can be reformulated as follows:

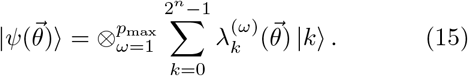

#### Ground-state calculation

By decomposing the Hamiltonian and state, leveraging the VQE algorithm, we can obtain an approximate ground state and its corresponding eigenvector of the large quantum system. From Equation (7), we aim to minimize the expectation value with parameter 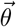,

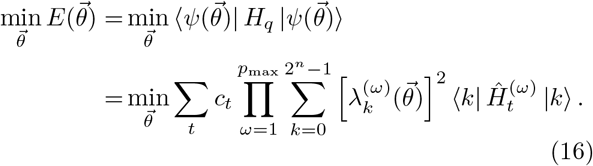

By utilizing the proposed approach, we can produce N-qubit quantum states and estimate the approximate ground-state energy of a 2^*N*^ -dimensional Hamiltonian using *p*_max_ *n*-qubit quantum processing units.

### 4.3 Problem-inspired ansatz

Because of the constraints of optimization problem, it offers an idea to exclude some infeasible answers and narrow the search spaces significantly, as well as speed up the convergence of the VQE algorithm for the hybrid *de novo* assembly problem.

Considering a specific value of *p*, the binary variable 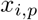 of Hamiltonian H_*q*_ can be encoded by state 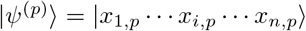. Because at most one node can be chosen for a position in the assembly path, one of binary variables x_*i,p*_ is limited to 1 or 0, while the remaining variables must be set to 0. After eliminating the incorrect solutions from the entire 2^*n*^-dimensional Hilbert space, the feasible answers are restricted to a set S of the n + 1 elements. Hence, it suffices to construct the following trial wavefunction:

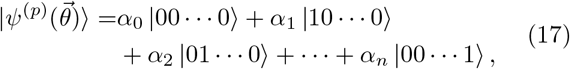

Where 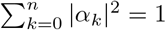.

Here, we construct the problem-inspired ansatz with *Rx, Ry* and CZ gates, shown in Figure 9, and apply it on the initial state |0 ⟩ ^*⊗n*^ to generate trial wave function 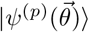 . To provide a more comprehensive understanding of the function of the quantum circuit, we have decomposed the process of preparing the trial wavefunction into two parts.

**Fig. 9.**
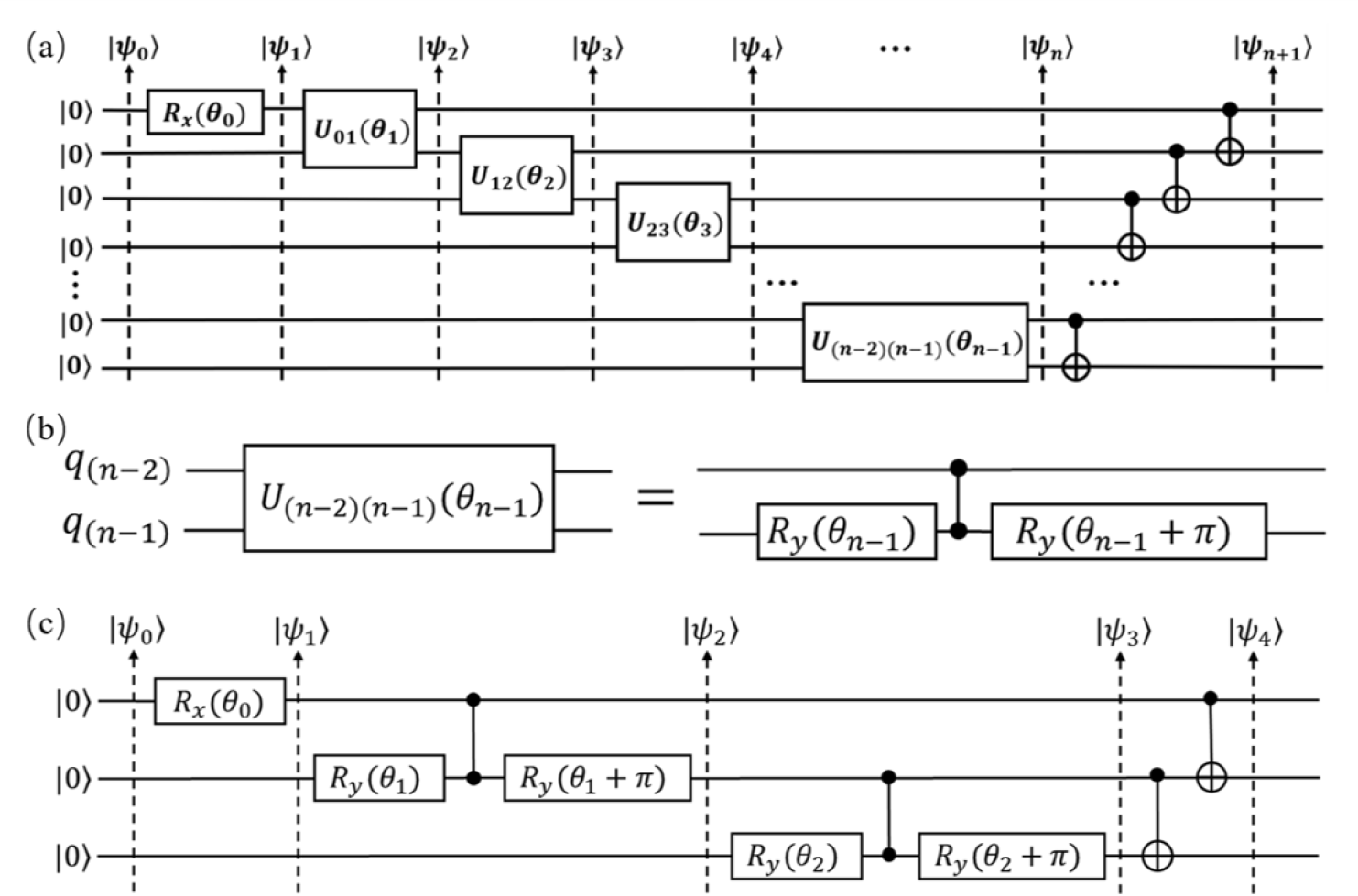
Schematic of the problem-inspired ansatz for preparing n-qubit wavefunction 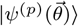. (a) Problem-inspired ansatz implemented by *R*_*x*_(*θ*_0_), *U*_*cq,tq*_(*θ*_*j*_) and CNOT gates. (b) Implementation of the *U*_*cq,tq*_(*θ*_*j*_) gate using *R*_*y*_(*θ*_*j*_), CZ and *R*_*y*_(*θ*_*j*_ +π) gates. (c) The quantum circuit for the 3-qubit wavefunction.

The first part involves the preparation of a pseudo-target state |ψ_*n*_ ⟩, which satisfies that the number of bases is equal to n + 1. In this part, our first step will be to initialize the state |ψ_0_ ⟩ = |0 ⟩ ^*⊗n*^. Then, a *R*_*x*_(*θ*_0_) gate is applied on the first qubit, thus giving us

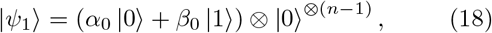

where α_0_ = cos (θ_0_/2) and β_0_ = −i sin (θ_0_/2).

Next, we apply a series of U_*cq,tq*_(θ_*j*_) gates in the next step, where *j* ∈ [1, n − 1]. As the Figure 9(b) shows, the U_*cq,tq*_(*θ*_*j*_) is implemented with R_*y*_(θ_*j*_), CZ_*cq,tq*_ and R_*y*_(θ_*j*_ + π) gates. Here, we set the cq-th qubit as a control qubit and the tq-th as a target qubit, respectively. When the state of the target qubit is |0⟩, the functions of U_*cq,tq*_(θ_*j*_) are as follows:

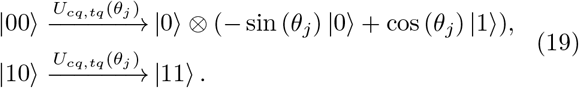

Thus, after a sets of *U*_*cq,tq*_(θ_*j*_) gates applied on |ψ_1_ ⟩, we obtain

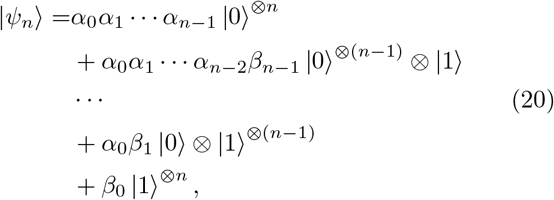

where α_*j*_ = − sin (θ_*j*_) and *β*_*j*_ = cos (θ_*j*_).

The second part involves a set of CNOT gates applied on pseudo-target state |ψ_*n*_⟩ to flip partial bases and prepare the trial wavefunction 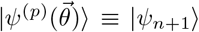. In order to better understand how a set of CNOT gates work, we consider a case with three qubits, and the quantum circuit is shown in Figure 9(c). The input state |ψ_0_ ⟩ = |000 ⟩ is sent through *R*_*x*_(*θ*_0_), *U*_01_(*θ*_1_) and *U*_12_(*θ*_2_) gates to give

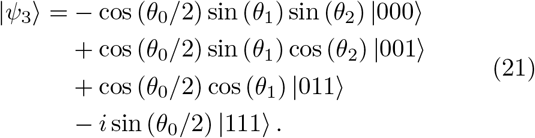

Compare to wavefunction |ψ_4_⟩, we need to flip the bases |011⟩ and |111⟩ in pseudo-target state |ψ_3_⟩ to |010⟩ and |100⟩, while keep others bases unchanged. Specifically,

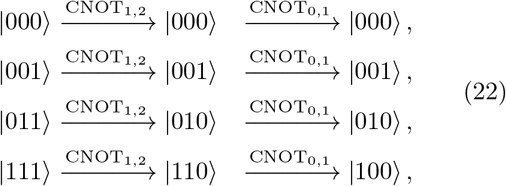

and

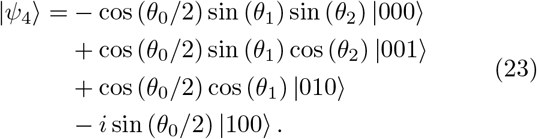

As shown in the example of three qubits, we can prepare 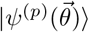 of any qubits by this problem-inspired ansatz. Here we give the matrix forms of the gates used in this ansatz. *R*_*x*_(*θ*_*j*_) and *R*_*y*_(*θ*_*j*_) are rotation gates acting on the j-th qubit. The CZ_*cq,tq*_ gate and CNOT_*cq,tq*_ are CZ gate and CNOT gate, respectively. And the unitary matrices of each gates are as follows:

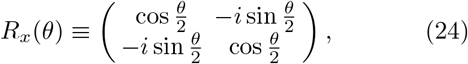

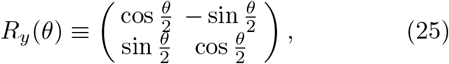

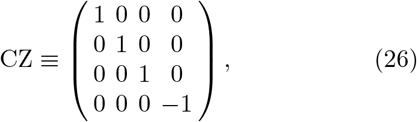

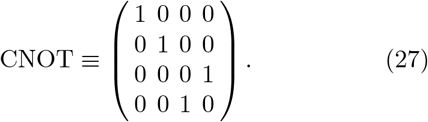

## Data availability

The data that support the findings of this study are available from the authors upon reasonable request.

## Code availability

The code that supports the findings of this study is available from the authors upon reasonable request.

## Acknowledgments

This work was supported by the National Key R&D Program of China (2022YFC3400400). We acknowledge support from China National GeneBank (CNGB) and the Guangdong Bigdata Engineering Technology Research Center for Life Sciences.

